# Tract-specific statistics based on diffusion-weighted probabilistic tractography

**DOI:** 10.1101/2021.03.05.434061

**Authors:** Andrew T. Reid, Julia A. Camilleri, Felix Hoffstaedter, Simon B. Eickhoff

## Abstract

Diffusion-weighted neuroimaging approaches provide rich evidence with which to estimate the structural integrity of white matter in vivo. Tract-based spatial statistics (TBSS), for instance, allows us to assess the spatial distribution of statistical tests performed on diffusion properties such as fractional anisotropy, which are proposed to reflect the integrity of white matter tracts. However, such methods do not provide a direct assessment of white matter integrity for connections between two specific regions of the brain. Here, we present a novel method for deriving tract-specific diffusion statistics, based upon arbitrarily-defined regions of interest. Our approach makes use of an empirically-derived population distribution based on probabilistic tractography, using the Nathan Kline Institute (NKI) Enhanced Rockland sample. We use a heuristic method to determine the most likely geometry of a path between two regions, if one exists, and express this as a spatial distribution. We then estimate the average orientation of streamlines traversing this path, at discrete distances along its trajectory, and compute the fraction of diffusion directed along this orientation for each participant. This allows us to obtain participant-wise metrics along the specific tract (tract-specific anisotropy; TSA), which can then be used to perform statistical analysis on any comparable population. Based on this method, we report both negative and positive associations between age and TSA for two networks derived from published meta-analytic studies (the “default mode” and “what-where” networks), along with more moderate sex differences and age-by-sex interactions in both networks.

## Introduction

Diffusion-weighted imaging (DWI) is a promising non-invasive in vivo technique for evaluating the integrity of myelinated axonal projections in the brain. DWI is based on the attenuation of the T2-weighted MRI signal in the presence of a field gradient, which indicates the degree to which diffusion is unrestricted in brain tissue in the direction of that gradient. Methods that reconstruct DWI maps across multiple gradient orientations can be used to model the diffusion of water molecules in discrete compartments (or voxels) of brain tissue, and the **anisotropy** of this diffusion can be used to estimate the orientation(s) along which diffusion is biased in that voxel. Because myelin is highly lipid-based, and forms a strong hydrophobic barrier, the degree of directed anisotropy in a white matter voxel is presumed to indicate the degree to which it is comprised of coherently oriented myelinated axons (***Assaf and Pasternak, 2007***). Moreover, variance in this anisotropy can be used to estimate the relative integrity of myelinated fibres that run through a voxel – with the assumption being that decreased anisotropy indicates a decrease in myelination or myelinated axons.

The simplest model of diffusion used for DWI analysis is the **diffusion tensor**, which assumes a Gaussian distribution with one principal and two secondary axes, corresponding to the first three eigenvectors of observed diffusion across gradient orientations (***Basser and Jones, 2002; Chung et al., 2010***). From this model, we can obtain a summary measure of anisotropy (fractional anisotropy; FA), based on the relative magnitudes of the eigenvalues associated with each axis of the tensor. FA ranges from 0 to 1, where 0 indicates perfect isotropy (as would be expected of a uniform substance such as water or cerebrospinal fluid), and 1 indicates diffusion exclusively in the principal direction. The diffusion tensor can also be used to perform **deterministic tractography**, in which streamlines are generated by starting at a pre-specified set of “seed” voxels, and propagated through a series of neighbouring voxels by reorienting at each according to its principal orientation of diffusion. Tractography can also be set in a probabilistic framework, by generating many streamlines and sampling at each voxel from a posterior probability distribution of orientations based on the tensor model (***Behrens et al., 2003***).

While the basic diffusion tensor is an adequate model for voxels through which fibres are essentially oriented in a single direction (e.g., fibres traversing the corpus callosum), it fails to model more complex situations, such as the case where two or more fibres are crossing, diverging, or converging. This presents a strong bias in favour of finding certain pathways over others. One way of addressing this issue is to explicitly model multiple fibre directions, for example by using Bayesian model estimation to determine whether multiple fibre orientations are present, and if so how strongly they contribute to the observed diffusion signal (***Behrens et al., 2007***). Such an approach greatly improves the ability of probabilistic tractography to discover tracts which traverse areas of uncertainty. As an example from Behrens et al. (2007): while seeding in the internal capsule led only to a prominent primary motor projection in the single-fibre approach, numerous other cortical targets were reached when crossing fibres were explicitly modelled.

It is often desirable to relate voxel-wise DWI-based metrics such as FA to other phenotypical observations, such as behavioural or cognitive measures, or clinical status. Voxel-wise analyses can be highly confounded by the individual geometry of white matter tracts, and one way to address this issue is tract-based spatial statistics (TBSS), in which FA measures are projected onto a population-based FA “skeleton” with a high probability of being white matter in all participants (***Smith et al., 2006, 2007***). The presence of crossing fibres, however, also has implications for the interpretation of FA (***Jbabdi et al., 2010***). As an example, for two identical fibres oriented along the anteroposterior (AP) axis, FA would be inversely proportional to the number of fibres crossing each along the (perpendicular) mediolateral (ML) axis. Interpreting FA in terms of the underlying microstructure of white matter in a voxel is thus inherently ambiguous. This ambiguity can be improved if crossing fibres are explicitly modelled, for example using the Bayesian approach described above. Such a crossing fibre model has been proposed as an extension to the TBSS approach (***Jbabdi et al., 2010***).

Although TBSS provides a means of assessing the spatial distribution of statistical effects on white matter integrity, it is often difficult to apply this distribution to specific connectivity in the brain. Suppose, for instance, that we are interested in whether the white matter comprising the physical connection between brain regions *R*_*a*_ and *R*_*b*_ is altered in condition *C*. Using TBSS, we observe that the *C*+ group has decreased FA in an area of white matter that *could* be intermediary to *R*_*a*_ and *R*_*b*_. However, as most major white matter tracts host a mixture of numerous projection, association, and commissural fibres, we can only really speculate about the possibility of a compromised *R*_*a*_*R*_*b*_ tract. To improve interpretability, we require an explicit approximation of the geometry of tract *R*_*a*_*R*_*b*_, and an estimate of the diffusion specifically oriented along this tract.

In this study, we introduce a novel methodology to address both of these issues. We first perform probabilistic tractography on a representative sample of participants (N=130, aged 18-80), using high angular resolution DWI data from the Nathan Klein Institute Enhanced Rockland sample (***Nooner et al., 2012***), and two sets of regions-of-interest (ROIs) obtained from published meta-analytic neuroimaging studies. For each pair of ROIs *R*_*a*_ and *R*_*b*_, we then compute the probability of a tract passing through a given voxel, and use a heuristic approach to determine whether a tract likely exists between *R*_*a*_ and *R*_*b*_, and what its most probable trajectory is. Finally, we use an approach similar to Behrens et al. (2007) to determine for each participant, and each voxel, the degree of diffusion in the direction of the tract at that voxel. This yields a tract-specific anistropy (TSA) metric that can be regressed against variables of interest. Here, we report the tract-wise and 3D distributions of TSA regressed against age, sex, and their interaction.

## Results

### Tract determination

We used a heuristic approach (see ***Figure 1***) to determine the most probable trajectory of a white matter projection between two regions of interest (ROIs). For a given pair of ROIs *R*_*a*_ and *R*_*b*_, we performed probabilistic tractography twice, seeding in one of these regions and terminating in the other. The resulting directed probability distributions were averaged to obtain a bidirectional probability average, which was then thresholded in order to identify a set of contiguous voxels connecting the two regions. If this step did not result in such a pathway, a connection between *R*_*a*_ and *R*_*b*_ was rejected; otherwise, we identified the “core” of the pathway and used this to obtain a final bidirectional tract trajectory estimate. Tract determination was carried out on two sets of ROIs, derived from previously published meta-analytic studies: the default mode network (DMN), and the what-where network (WWN).

**Figure 1.**
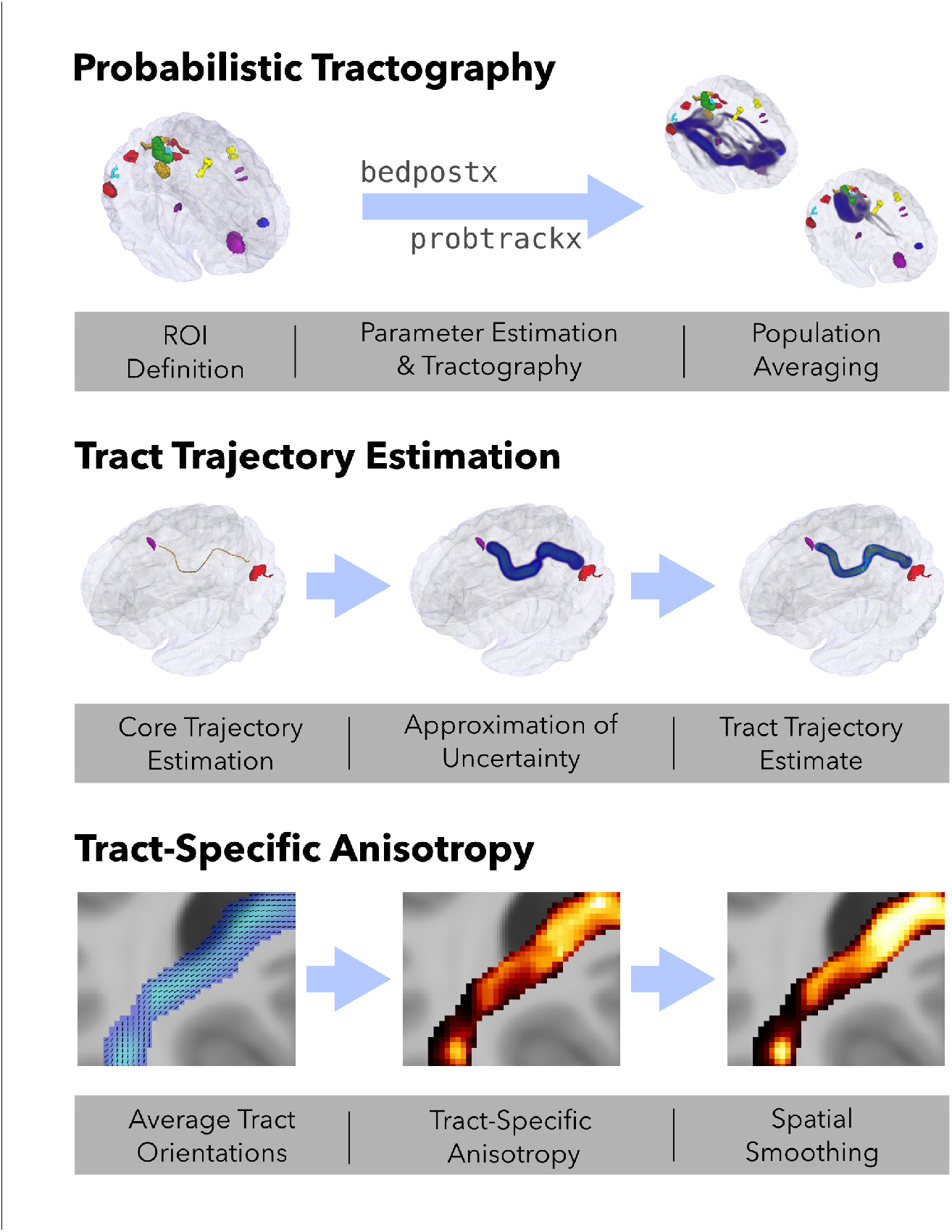
Schematic of the procedure followed in this study. *Probabilistic tractography*: ROIs were obtained from previous meta-analytic studies and used as seed/target regions for diffusion tensor modelling with `bedpostx` and probabilistic tract tracing performed with `probtrackx`, across all participants. The resulting probability distributions were averaged across directions for each ROI pair (**P**_*ab*_). *Tract trajectory estimation*: For each ROI pair, a “core” trajectory was estimated from these bidirectional averages, and represented as a 3-dimensional polyline. An uncertainty field (Ф_*ab*_) was then generated from this polyline using an anisotropic Gaussian kernel. Finally, a “core” tract estimate (**P**_*ab*−tract_) was generated as the element-wise product of **P**_*ab*_ and **Ф**_*ab*_. *Tract-specific anisotropy*: Average tract orientations were computed for each voxel in a given tract, and these orientations were then regressed against the diffusion evidence for each individual participant. This produced a tract-specific anisotropy (TSA) distribution for each participant, that can be regressed against variables of interest.

***Figure 2***a illustrates the tract trajectory estimation process for example tracts PFCm(R) - LOC(R) and dPMC(L)-vPMC(R). The horizontal and coronal slice renderings show the initial minimal bidirectional averages (top row) and final tract trajectory estimates (bottom row). Figure 2b shows three-dimensional renderings of these steps, including the failure to estimate a core trajectory for SPL(L)-dPMC(R). For the vast majority of tracts, this method was able to isolate a single, core trajectory from a variety of alternatives.

**Figure 2.**
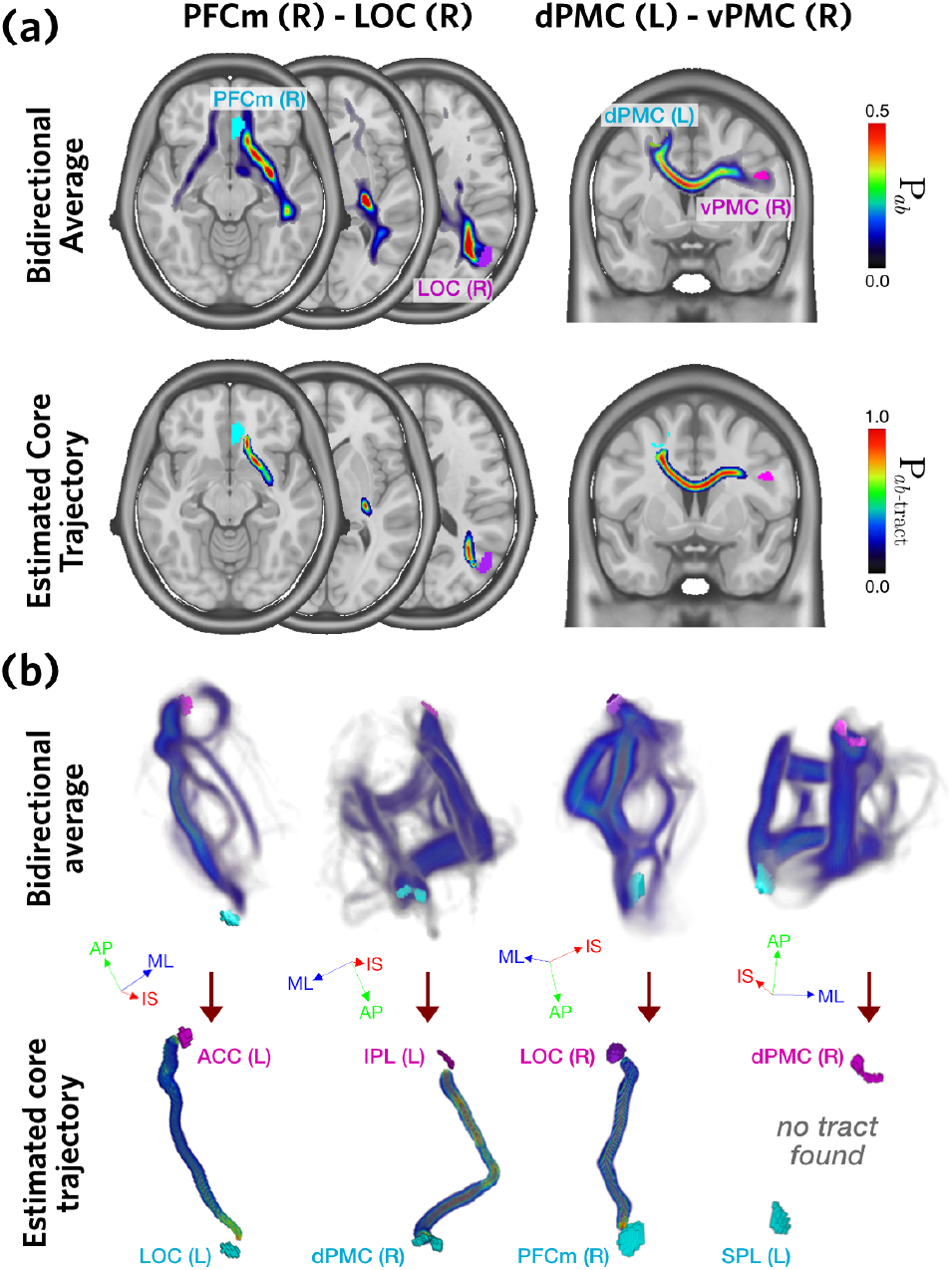
**(a)** Core tract trajectory estimation steps, shown for two exemplar tracts. The top row shows the streamline probability determined across all participants, averaged in both directions, **P**_*ab*_. The bottom row shows the estimated core trajectory for each ROI pair, **P**_*ab*−tract_. **(b)** Three-dimensional distributions of **P**_*ab*_ (top) and **P**_*ab*−tract_ (bottom), for four exemplar tracts. Axis labels: ML, medial-lateral; AP, anterior-posterior; IS, inferior-superior.

For the DMN, 33 of 36 ROI pairs (92%) produced core tract estimates (***Figure 3***). Two of the failed tracts involved the left PFCm, with the more posterior LOC(R) and PCm(L) regions. Tracts connecting PCm and PCG to PFC and ACC traversed the more superior cingulum bundle, while those connecting LOC to PFC and ACC traversed the more inferior fronto-occipital fasciculus. Contralateral DMN connections traversed the splenium of the corpus callosum (CC; Figure 3d) for posterior ROIs, and the genu of the CC for anterior ROIs and anteriorly projecting LOC connections.

**Figure 3.**
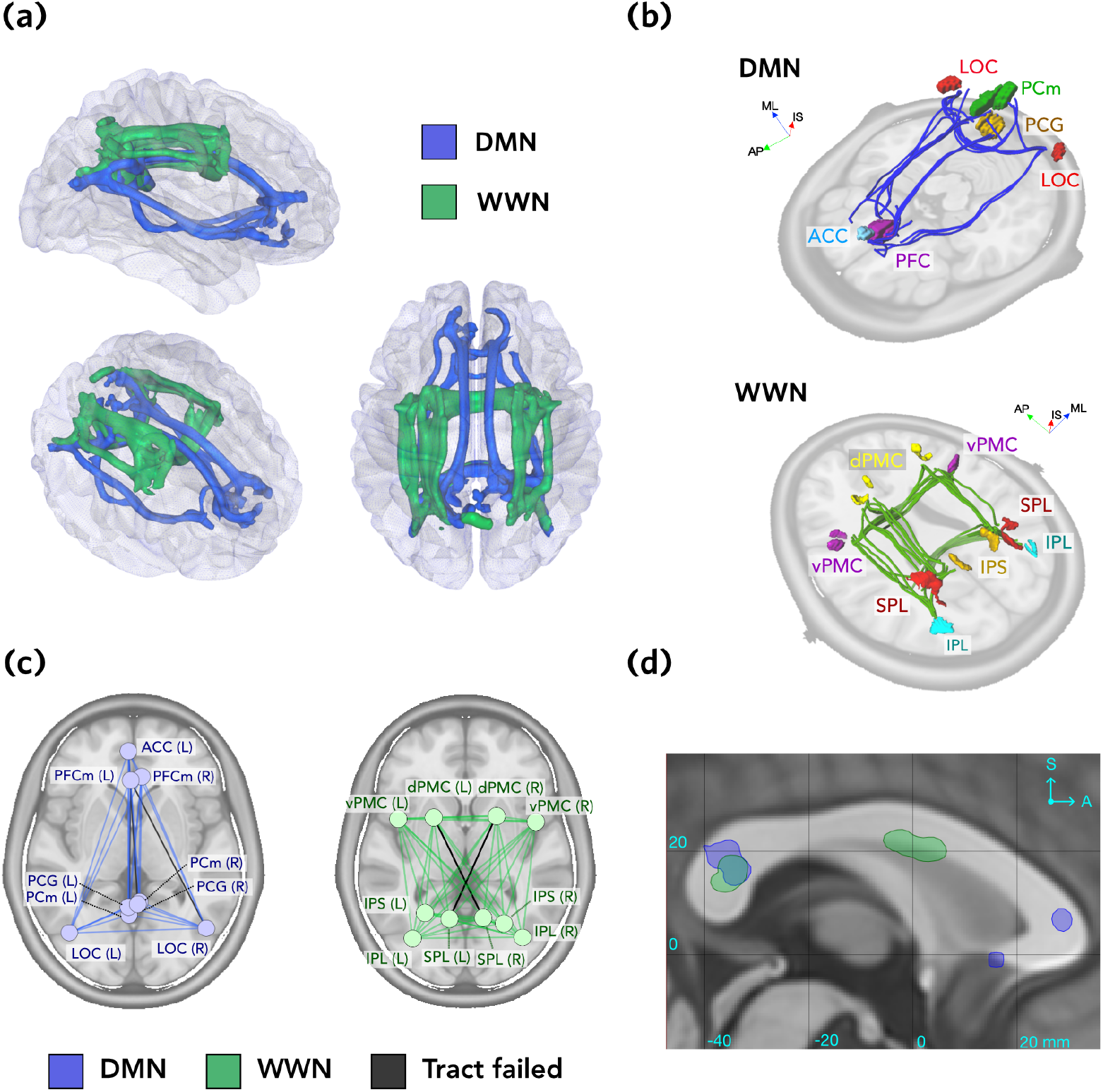
Tract determination. **(a)** All accepted tracts shown for the DMN (blue) and WWN (green) networks, rendered as isosurfaces thresholded at 0.5 (top, right, and oblique perspectives). **(b)** Core polylines representing the trajectories of accepted tracts in the DMN (top) and WWN (bottom) networks. Images are rendered with oblique perspectives next to planar sections of the ICBM non-linear T1 template image, for reference. See Materials and Methods for names of the ROIs. **(c)** Graph representations of the DMN (blue) and WWN (green) networks, with failed edges shown in grey. Graph vertices represent the center points of each ROI, projected onto the transverse plane. **(d)** Sagittal section at midline, showing where contralateral tracts (thresholded at 0.5) for each network traverse the corpus callosum. Coordinates are ICBM152. Axis labels: ML, medial-lateral; AP, anterior-posterior; IS, inferior-superior.

For the WWN, 43 of 45 tracts were estimated (96%), with only 2 tracts failing because no core tract could be identified (***Figure 3***). The failed tracts were the two contralateral connections between SPL and dPMC (Figure 3c). Successfully estimated tracts consisted of anteroposterior projections traversing the superior longitudinal fasciculus, and contralateral connections traversing the splenium of the CC and the inferior part of the body of the CC (Figure 3d).

### Tract-specific anisotropy

Given a tract trajectory estimate, as described above, we next wanted to determine how strongly diffusion profiles of individual participants were oriented along that tract. For a given voxel, we determined the average streamline orientation derived from the previous probabilistic tractography step, and then computed, for each participant, a “tract-specific anisotropy” (TSA) estimate, indicating the degree to which this average orientation loaded onto the DWI intensities along each gradient direction (TSA values are the β parameters estimated from ***Equation 3***).

Kernel density estimates for TSA, for both networks, are shown in ***Figure 4***a. These show the distribution of scores for tracts thresholded at **P**_*ab*−tract_ > 0.5. While most distributions were roughly Gaussian, with a heavy positive tail, this varied across tracts in terms of both kurtosis and skewness, and some shorter tracts (e.g., PCG(L)-PCm(R)) had very little variance. For a number of tracts (e.g., SPL(R)-IPS(L)), a bimodal distribution was evident. The overall distribution for each network is shown in the insets; WWN showed a broader, flattened distribution relative to DMN.

**Figure 4.**
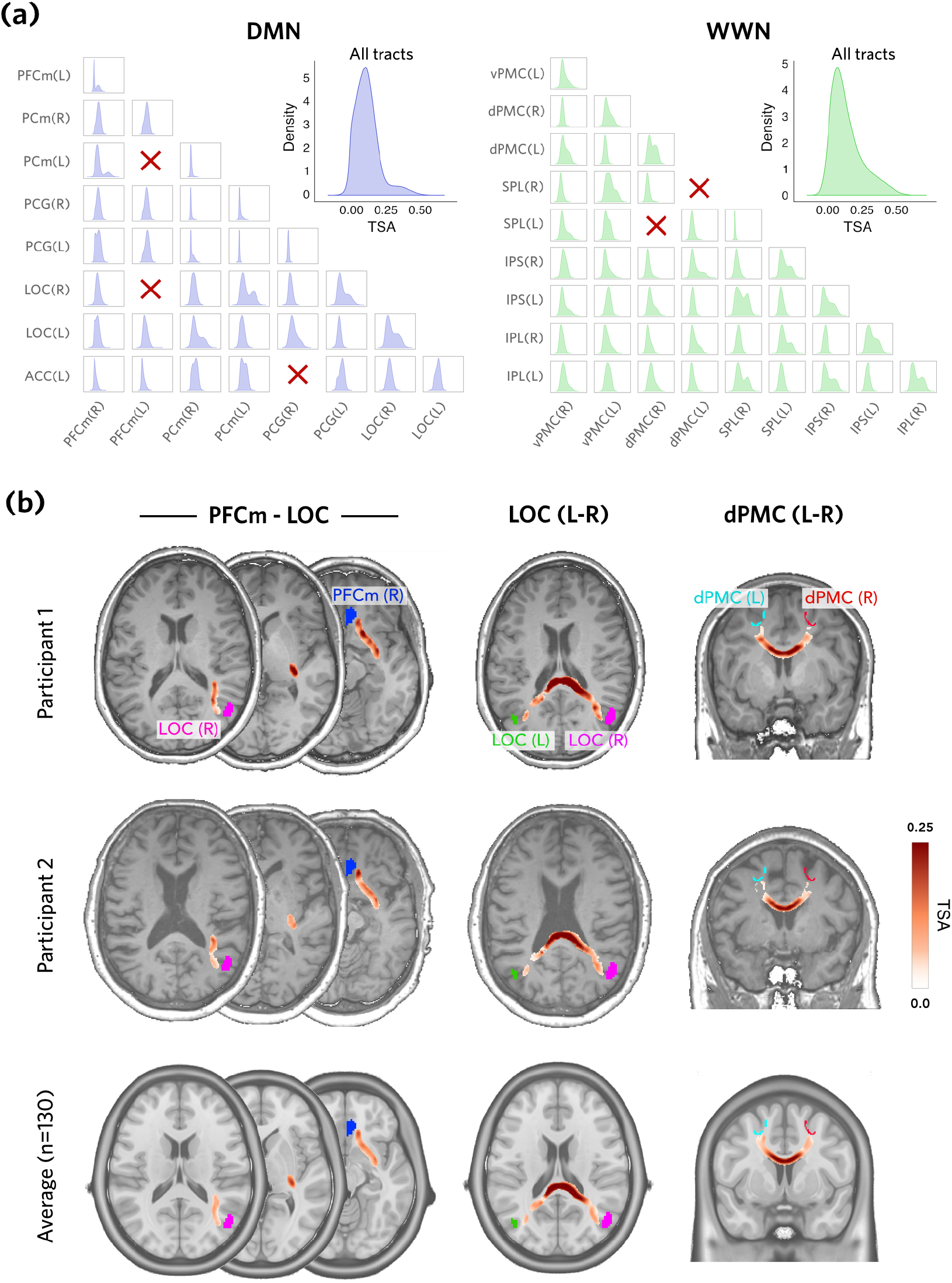
Tract-specific anisotropy (TSA). **(a)** Distributions of TSA across voxels and participants, for each estimated tract (ROI pair), shown as kernel density estimates for DMN (blue) and WWN (green). Insets show the distribution of TSA values over all tracts; x-axis scales are the same for all plots. Red crosses indicate that no tract was generated. **(b)** Spatial distributions of TSA values for two exemplar participants and averaged over all participants, and three examplar tracts, shown in horizontal and coronal section. Anatomical images are the ICBM152 nonlinear template.

***Figure 4***b shows the two-dimensional spatial distributions of TSA scores for two exemplar participants, along with the average TSA across all 130 participants. These distributions demonstrate variability between participants, but in general the highest TSA scores were observed in the center of estimated tract trajectories - reflecting the strongest loading of diffusion profiles - which tapered off towards the edges.

### Tract-specific Age and Sex effects

Having obtained TSA scores for each individual participant, we were next interested in whether these scores were associated with age, sex, and their interaction. We performed voxel-wise regressions of the form *TSA* = *β*_0_ + *β*_1_ ·Age + *β*_2_ ·Sex + *β*_3_ ·Age × Sex +*ε*. T-statistics for each coefficient were obtained, and summarised at each distance along the tract. For each tract, we then used one-dimensional random field theory to identify significant clusters of t-statistics (p<0.05). To control for family-wise error, we limited the false discovery rate (FDR) over all tracts to 0.05.

For the DMN, there were significant associations of TSA with age and sex, as well as the interaction between these two factors (***Figure 5***, top row). Age effects were fairly diffuse. Negative effects of age were found in 15/32 (47%) of DMN tracts, including ipsi- and contralateral connections between PFC and LOC (traversing the fronto-occipital fasciculus), and PCm(L)-PFCm(R). Modest positive effects of age were also found for 4/32 (12 %) of tracts. These were smaller, and comprised exclusively of ipsilateral tracts (three left and one right). Both positive (female > male) and negative sex differences were also found. ***Figure 6*** shows scatter and violin plots for selected tracts. Notably, tract PCm(L)-PCG(L) showed effects for age, sex, and their interaction; and tract ACC(L)-LOC(L) and PFCm(R)-LOC(R) showed both positive and negative age effects.

**Figure 5.**
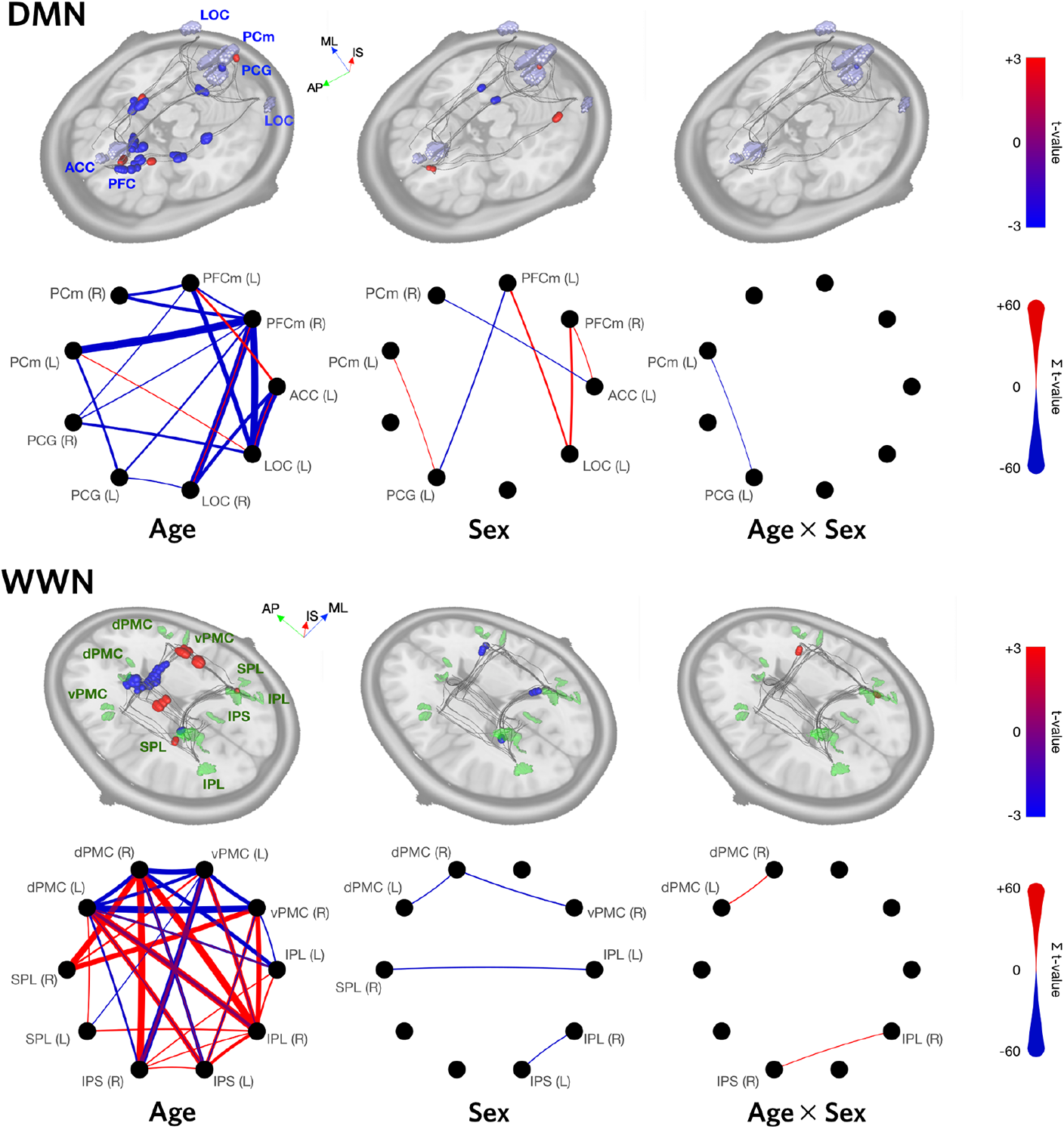
Regression results for both networks, for linear models of the form *TSA* = *Age* + *Sex* + *Age* × *Sex* + *ϵ*. 3D renderings show the maximal t-values (thresholded using cluster-wise inference, with FDR<0.05) at each distance along the core trajectories of each tract in the network. Circular graph representations show the sum of significant positive (red) and negative (blue) t-values for each tract. The thickness of an edge is proportional to its sum, and the absence of an edge indicates that no significant clusters were found for that tract. Axis labels: ML, medial-lateral; AP, anterior-posterior; IS, inferior-superior **Figure 5–Figure supplement 1**. GLM t-value distance traces for all ROI pairs in the DMN. **Figure 5–Figure supplement 2**. GLM t-value distance traces for all ROI pairs in the WWN.

**Figure 6.**
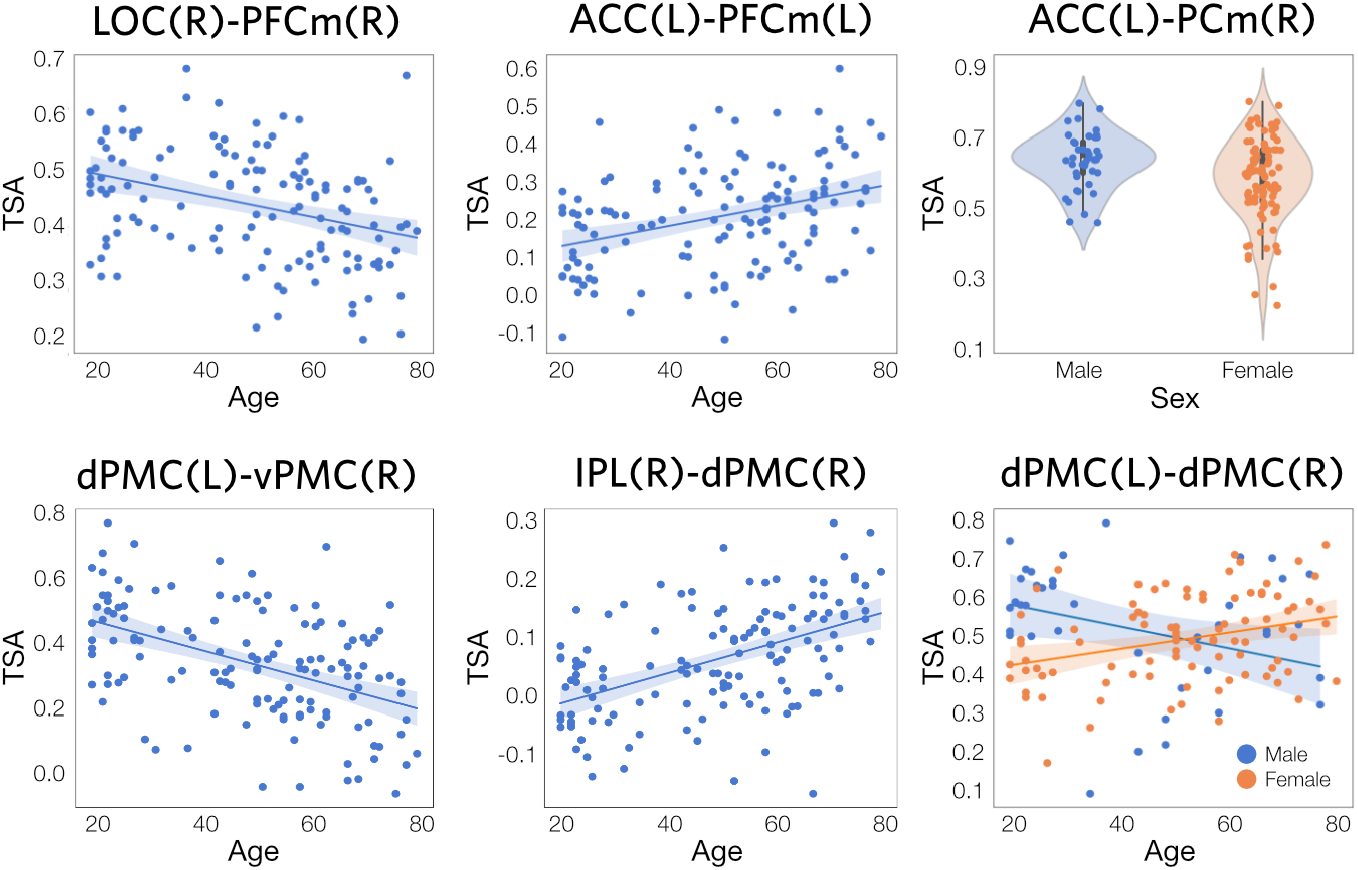
Plots of selected TSA regression results. *Top row*: DMN tracts with (from left to right) negative and positive age effects, and a sex difference (male > female). *Bottom row*: WWN tracts with negative and positive age effects, and an Age×Sex interaction. Data points are the mean TSA values within significant clusters (as shown in ***Figure 5***). Shaded areas on scatterplots represent 95% confidence intervals. **Figure 6–Figure supplement 1**. Scatter and violin plots for all significant effects in the DMN. **Figure 6–Figure supplement 2**. Scatter and violin plots for all significant effects in the WWN (Part A). **Figure 6–Figure supplement 3**. Scatter plots for all significant effects in the WWN (Part B).

The WWN also showed significant associations for age, sex, and their interaction (*Figure 5*, bottom row). Surprisingly, there were strong posirtive and negative age effects for this network, with 16 /32 (50%) having showiung negsative effects and 14/32 tracts (44%) showing positive effects. Compared to DMN, the age effects for WWN were more focal. Negative age effects were found proximal to dPMC(L), and in collosal fibres traversing the body of the corpus callosum (see ***Figure 3***d), involving contralateral tracts between dPMC and vPMC. There were also negative associations found in contralateral tracts between IPS, dPMC, and vPMC. The largest positive age effects occurred along anteroposterior-oriented tracts in the longitudinal fasciculus, particularly those involving left and right dPMC. All sex differences were negative (male > female) for this network, on were found on contralateral tracts. ***Figure 6*** shows scatter and violin plots for selected tracts. Notably, the homotopic dPMC(L)-dPMC(R) tract showed both a sex difference and an AgexSex interaction.

Scatter and violin plots for all significant effects, and line plots and tables showing raw and cluster-thresholded t-values for all tracts, are provided in Supplementary Material.

## Discussion

### Estimation of core tract trajectories

In this study we introduce a novel method for obtaining tract-specific spatial estimates of the “core” trajectories of projections between two arbitrarily defined ROIs, on the basis of probabilistic DWI tractography performed across a representative cohort of healthy individuals. This approach yielded plausible trajectory estimates for 33 of 36 ROI pairs (92%) in the DMN network, and 43 of 45 ROI pairs (96%) in the WWN network. In order to achieve sufficient coverage of the space of possible trajectories, we derived our probability estimates from a large number of streamlines (50,000 × ROI) over 130 participants. Our results demonstrate the feasibility of using DWI-based probabilistic tractography for delineating individual tracts between arbitrary pairs of distal brain regions, as well as obtaining participant- and tract-specific anisotropy values from these tracts.

ROI pairs failed to generate tract trajectory estimates when the thresholding applied to the bidirectional population average distribution broke the contiguous path between them. Notably, this is not evidence that the tract does not exist, but rather that there is ambiguity about the location of its trajectory, based on the diffusion evidence. It is important to note that the threshold applied here was determined somewhat arbitrarily, i.e., by observing what value reduced the number of alternative pathways to a single, core one. In cases where this failed, there were typically two or more alternatives that could not be disambiguated (see ***Figure 2***b for an illustration of failed tract SPL(L)-dPMC(R)). Further investigation into these ROI pairs, e.g., by comparing them to known connectivity evidence from tract tracing studies (in non-human primates) or different modalities (in humans), may be useful for determining whether a tract indeed exists between them.

Previous studies have also evaluated white matter tract geometry using DWI. In a now-classic study, ***Catani et al. (2002***) used deterministic tractography to identify and “dissect” individual tracts based on seed ROIs. This was extended by ***Jones et al***. (*2005*) in order to map diffusion metrics onto the trajectories of specific tracts. More recently, ***Colby et al. (2012***) used B-spline resampling to map FA to specific tracts in individual participants, allowing effects to be mapped to points along its trajectory. The present approach extends the work of these previous studies, in that it facilitates the investigation of white matter pathways connecting specific ROI pairs, rather than coarse-scale fasciculi, in terms of tract geometry and anisotropy estimation. Another important advantage is its use of a probabilistic tractography framework to describe the “core” trajectories of specific tracts across a population of interest.

### Tract-specific anisotropy

It is common to use streamline counts from probabilistic tractography as an estimate of connection strength between two ROIs (e.g., ***Daianu et al., 2013; Lohse et al., 2014; Reid et al., 2016b***). This approach, however, suffers from a number of seemingly intractable biases, including: the nature of the diffusion profile through which a given tract traverses (anisotropy bias); the length of the tract (distance bias); and the position of seed and target ROIs relative to gryi or sulci (***Reveley et al., 2015; Reid, 2016***). It is thus uncertain how to interpret a relative streamline count, or change in this quantity in association with some covariate of interest, with respect to biophysical properties such as white matter integrity or connectivity strength. The present approach is arguably less susceptible to these biases, because it only models the geometry of streamlines that *do* connect two ROIs. However, future research efforts should be directed at validating (1) the interpretability of TSA with respect to axonal white matter integrity, and (2) the spatial specificity of trajectory estimates. This could be done, for example, through the use of phantoms (***Zhu et al., 2011; Caspers and Axer, 2017***) or histological approaches (***Mollink et al., 2017; Schaeffer et al., 2018***).

The present method might be considered an extension of the well-known TBSS approach (***Smith et al., 2007***), in which statistics are projected onto a pre-established population-based white matter “skeleton”, which serves a similar function to population-based anatomical grey matter templates, such as the linear and nonlinear ICBM-152 templates (***Fonov et al., 2009; Mazziotta et al., 1995***). There are two major advantages of the present method over TBSS: (1) it allows a population-based tract estimate to be *derived specifically for the white matter tract connecting any two arbitrarily-defined ROIs*, if that tract is likely to exist; and (2) it *allows participant- and tract-specific anisotropy to be estimated*, based on the orientation of streamlines defining that tract in each voxel along its trajectory. Conversely, one major limitation with the TSA approach is that it is substantially more computationally intensive, and scales quadratically with the number of ROIs in a network of interest.

### Age and sex are associated with tract-specific anisotropy

Positive and negative age/TSA associations were found for both networks. There is an abundance of evidence for age-related decreases in DMN connectivity, including a decrease in fMRI-based resting-state functional covariance (***Hafkemeijer et al., 2012; Marstaller et al., 2015; Sambataro et al., 2010***), and age-related decreases in FA in white matter tracts proximal to DMN regions (***Marstaller et al., 2015***). Marstaller and colleagues found a reduction in the extent to which posterior cingulate and precuneus activity covaried with wider brain networks, along with reduced FA in numerous tracts, including the fronto-occipital fasciculus, where the current negative age association is most prominent. Sambataro et al. reported decreased covariance between PCC and PFC, which predicted working memory performance. Decreases in DMN functional covariance also appear to be accelerated in Alzheimer’s disease (***Jones et al., 2011***; also reviewed in ***Reid and Evans, 2013***).

For the WWN, age-related decreases in TSA occurred mainly in the body, but not the splenium, of the corpus callosum. This pattern is in agreement with several DWI-based studies of age-related connectivity changes. ***Burzynska et al. (2010***) used TBSS to show an age-related reduction in FA (and increase in radial and mean diffusivity) in the genu and body of the corpus callosum, but not the splenium. Using a DTI approach, ***Bennett and Madden (2014***) found decreased FA in older versus younger participants for both the genu and splenium, but with a more pronounced effect in the former. The same authors report an anterior-to-posterior gradient in age-related FA changes, with these being more pronounced in frontal white matter, consistent with the pattern found in the current study (***Bennett et al., 2009***).

Positive age associations were a more surprising finding, as numerous articles report negative age/FA associations (e.g., ***Kodiweera et al., 2016***) and postmortem evidence of white matter loss and decrease in the proportion of small myelinated fibres with age (***Tang et al., 1997; Aboitiz et al., 1997***). Both positive and negative age/FA associations have been reported in at least one previous brain-wide TBSS study (***Kochunov et al., 2007***), however. For DMN in particular, we found that most positive associations occurred in regions proximal to ROIs, where the potential confound of crossing fibres is likely more pronounced (***Jeurissen et al., 2012***). The positive relationships in the WWN were especially prominent, and suggest a (paradoxical) increase in white matter integrity in these tracts. The majority of positive relationships in WWN were found in the middle of the superior longitudinal fasciculus (SLF). One TBSS study focusing on the SLF in a healthy cohort found no effect of age on FA (***Madhavan et al., 2014***), while another whole-brain study found SLF among tracts with negative age effects (***Marstaller et al., 2015***). Increased FA in Alzheimer’s disease patients has also been reported in SLF (***Douaud et al., 2011***). The authors of this study suggest that a relative sparing of crossing motor fibres may account for this effect, but this is consistent with our observed increase in TSA; on the contrary, our findings might be explained by a relative *decrease* in WM integrity in these crossing fibres.

It is possible that the increased specificity of the current approach permits a more fine-grained spatial and angular dissection of effects than does TBSS with FA, which uses a more coarse-grain white matter skeleton, and is not orientation-specific. If so, then the positive effects observed here may reflect a real age-related increase in white matter integrity for specific tracts. This possibility is supported by reports of fairly widespread increases in fMRI-based functional covariance with aging (***Tomasi and Volkow, 2011***), which have been proposed to reflect compensatory changes in response to degeneration or dysfunction of other brain regions. Given the conflicting evidence, however, these effects should be interpreted with caution. It will be important in future TSA studies to increase the number of ROIs, or query specific crossing tracts, in order to obtain a more complete picture of age-related effects across white matter.

More modest sex differences, and age-by-sex interactions, were also observed for both networks. Previous DWI-based studies have also found sex differences in FA, typically with males showing higher FA values (***Takao et al., 2013; Rathee et al., 2016; Menzler et al., 2011***). For DMN, we found negative effects (males > females) for the PCG(L) with PFCm(L), which accords with findings from ***Menzler et al. (2011***), who report prominent differences in the cingulum bundle. We also found positive effects (females > males) that were mostly left lateralized and included LOC(L), which have not commonly been reported in the literature. Although sex differences in LOC function have been hypothesised on the basis of differential object perception (***Vanston and Strother, 2016***) – a function that has been associated with this region – there does not currently appear to be much direct evidence for this.

It is also notable that, for numerous tracts, we found both positive and negative age effects in different parts of the same tract (see ***Figure 5***). It is not immediately clear how to interpret such a result. If our inference is that a TSA value represents the number of intact axons projecting between two grey matter ROIs, this is contradicted by the observation of both increases and decreases in this region - damage to an axon anywhere along its length will trigger Wallerian degeneration in both directions. On the other hand, this finding is consistent with the idea that TSA reflects the degree of (de)myelination, which may be increased on average in one part of the tract and decreased in another. Importantly, there is evidence that demyelinated axons, while they may be functionally impaired, are not necessarily at higher risk of degeneration (***Smith et al., 2013***). A third possibility is that TSA is influenced by the degree of crossing fibres in a particular region along its length; changes to diffusion in directions other than the average orientation of interest will influence the regression fit used to estimate this metric.

### Limitations and future directions

The NKI Rockland dataset was chosen due to its large size, age range, and the use of a single MRI scanner and protocol. To ensure the cohort was as representative of the general population as possible, and to enable the analysis of age over the lifespan, we chose to use close to the full age range (18-80), and to exclude participants with clinical diagnoses. As with most population templates, however, the choice of cohort is an important consideration when interpreting a derived result. The human brain is known to show systematic anatomical grey matter changes across the lifespan (***Sowell et al., 2003; Giorgio et al., 2010***), and this will almost certainly bias normalization in a way that may account for a portion of the TSA effects reported here. It will be important in future studies to assess the influence of this bias, use cohorts that are more targeted to a particular phenomenon under investigation, and compare the predictions of TSA to in vivo or post mortem analyses of white matter (e.g., as in ***Reveley et al.,2015***).

The ability to estimate spatial trajectories of white matter projections between specific pairs of ROIs could lead to a number of important applications. For instance, this information could be used to predict the functional outcomes of age-related white matter lesions (WML), which have a prevalence of ∼18% in people aged 60-69 and ∼40% in people over 80 (***Vernooij et al., 2008***). Conceivably, clinicians would identify WML locations using T2-FLAIR imaging, estimate which tracts intersect these lesions, and predict functional deficits on the basis of the brain regions connected by these tracts. This approach could augment previous studies investigating connectivity (***Tuladhar et al., 2014***), cortical thickness (***Reid et al., 2010***), and gait changes related to WML (***de Laat et al., 2012***).

## Materials and Methods

### Participants

Participant data were obtained from the publicly available Nathan Klein Institute (NKI) Enhanced Rockland sample (**Nooner et al**., **2012**), through the 1000 Functional Connectomes Project (www.nitrc.org/projects/fcon_1000/). We included participants from the first four data releases, while excluding any participant with an existing clinical diagnosis at the time of scanning. In total, 130 participants (86 female, age range 18 to 80) were analyzed.

### Neuroimaging data and metadata

All imaging data in the Rockland sample were acquired from the same scanner (Siemens Magnetom TrioTim, 3.0T). T1-weighted images were obtained using a MPRAGE sequence (TR = 1900 ms; TE = 2.52 ms; voxel size = 1 mm isotropic). DWI was collected with a high spatial and angular resolution (TR = 2400 ms; TE = 85 ms; voxel size = 2 mm isotropic; b = 1500 s/mm2; 137 gradient directions). Age and sex data were also obtained, with the Sex factor encoded as 1=Male, 2=Female.

### Preprocessing

DWI images for all participants were preprocessed using FMRIB software (***Smith et al., 2004; Woolrich et al., 2009***); specifically, FSL version 6.0.0. Participant datasets were processed in parallel using a Linux-based SLURM computing cluster, located at the University of Nottingham. Raw diffusion data were first corrected for eddy current artifacts using the eddy command. All *B*_0_ (zero-gradient) images were then averaged and used to extract a brain mask using the bet command. Next, for each voxel in the brain mask, a diffusion tensor model was fit to the data using dtifit, and subsequently passed to the bedpostx command, which infers the existence of crossing fibres and estimates the contribution of each crossing fibre to the diffusion-weighted signal (***Behrens et al., 2007***). Finally, to express the voxel-wise diffusion models in standard space, linear and nonlinear transforms between native diffusion and MNI-152 space were estimated, by applying the `flirt` (12 degrees of freedom) and `fnirt` commands, respectively, using individual FA images as input and a mean template FA image as the reference. Inverse transforms were also estimated, using the invwarp command, for use in later steps.

### Regions of interest

Seed regions of interest (ROIs) were obtained from two previously-defined networks (see ***Figure 3***b). For each network, all possible ROI pairs were considered as potential tracts.

The *default-mode network* (DMN) was derived from ***Schilbach et al. (2012***), who performed an Activation Likelihood Estimation (ALE; ***Eickhoff et al., 2009, 2012***) meta-analysis using 533 experiments from the BrainMap database (***Fox and Lancaster, 2002***), querying for regions that were consistently deactivated across tasks. The ROIs were obtained by downloading the DMN network from the ANIMA database (https://anima.inm7.de/studies/Schilbach_SocialNetworks_2012; ***Reid et al., 2016a***). The network was comprised of ROIs: anterior cingulate cortex (ACC; left only), lateral occipital cortex (LOC; bilateral), posterior cingulate gyrus (PCG; bilateral), precuneus (PCm; bilateral), and medial prefrontal cortex (PFCm; bilateral).

The *what/where* network (WWN) was derived from ***Rottschy et al. (2012***), with data obtained from the ANIMA database (https://anima.inm7.de/studies/Rottschy_WorkingMemory_2012). The authors performed a conjunction on ALE analyses for tasks testing memory for object identity (“what”; 42 experiments) and object location (“where”; 13 experiments). The WWN was comprised of 10 ROIs, with 5 brain regions represented bilaterally. These ROIs were: dorsal and ventral premotor cortex (dPMC, vPMC), superior parietal lobule (SPL), inferior parietal lobule (IPL), and infraparietal sulcus (IPS).

### Probabilistic tractography

Probabilistic tractography was performed in MNI-152 space, using the `probtrackx` command (***Behrens et al., 2007***). For both networks, we generated 50,000 streamlines per seed voxel, with the other seed regions as target masks. We applied a step length of 0.5 mm, a curvature threshold of 0.2, a minimal path distance of 5 mm, and a fibre threshold of 0.01. A distance correction was also applied. For each voxel `probtrackx` recorded the number of streamlines which encountered that voxel, producing a separate count for streamlines terminating in each target ROI. Additionally, the voxel-wise mean orientation of streamlines between each seed/target pair was computed.

### Tract determination

Tracts were determined separately for each ROI pair. Voxel-wise streamline counts for each direction (*R*_*a*_ to *R*_*b*_ or *R*_*b*_ to *R*_*a*_) were first normalized by dividing by the total number of streamlines reaching target *R*_*b*_ from seed *R*_*a*_, and the minimum across both directions was obtained. These minimum images were subsequently averaged over participants, smoothed, and normalized to the range [0,1], yielding a probability *P*_*ab*_(*i*) of voxel being included in tract *T*_*ab*_. It is noteworthy that in some cases, the matrix **P**_*ab*_ of such values is constrained to a single, geometrically confined tract, while in others, it reflects many possible routes between two ROIs (see examples in ***Figure 2***b). Our objective was to identify the single most probable tract, or reject the tract altogether if such could not be determined. This was done in a heuristic manner. Firstly, we thresholded **P**_*ab*_ by setting all values where **P**_*ab*_ <*α* to zero. A threshold of *α* = 0.07 was applied by experimentation, in order to both disconnect most low-probability alternative pathways, and ensure that (for cases where the existence of a tract is known or likely) at least one pathway remained intact. Additionally, because **P**_*ab*_ proximal to ROIs tended to be lower that for the main tract trajectory, ROIs were dilated by 3 voxels to ensure they remained connected after thresholding.

We next discretized **P**_*ab*_ into distance assignments **D**, using a flood-fill approach, assigning all voxel neighbours of seed region ***R***_*a*_ a value of *d*= 1, all subsequent neighbours *d*= 2, and so on, until all voxels in *T*_*ab*_ were assigned a distance. We then identified the voxel of maximal *P*_*ab*_ for each distance, constructing a polyline **L**_*ab*_ with center points of these voxels as its vertices. Inclusion of a vertex in **L**_*ab*_ was conditional on two constraints: (1) segment length |**x** _*i*−1,*i*_|< 4 mm; and (2) vertex angle *θ*_*i*_ < *π*/3. Where a constraint was violated, the voxel with the next highest *P*_*ab*_ was tested, and so on until a voxel was found satisfying the constraints. In cases where no appropriate voxels were found, the maximal voxel was added and the violation was recorded as a flag for visual inspection. **L**_*ab*_ was extended in this fashion until: (1) the target region *R*_*b*_ was encountered, and the tract was accepted for further analysis; or (2) the maximal distance was encountered, but not the target region *R*_*b*_, and the tract was rejected for further analysis (i.e., the thresholding broke all routes, and thus a “true” route between *R*_*a*_ and *R*_*b*_ could not be determined).

With the assumption that an accepted accepted polyline represents the geometric center of a given tract *T*_*ab*_, we then modelled the probability **P**_*ab*−tract_ of a voxel being in that tract, as the product of an uncertainty field **Φ**_*ab*_ oriented around **L**_*ab*_, and the original probability field **P**_*ab*_ (see *Figure 1*).

**Φ**_*ab*_ was constructed as follows. For every vertex *v* ∈ **L**_*ab*_, an orientation vector *ω*_*v*_ was computed as the sum of the segment vectors **x**_*v*−1,*v*_ and **x**_*v,v*+1_ (at the endpoints of **L**_*ab*_ only one segment was used).From *ω*_*v*_, an anisotropic Gaussian kernel *ϕ* (*i,ω*_*v*_, *µ, σ*_*a*_, *σ*_*r*_) was applied to the subset of voxels within a radius of 8 mm of *v*. Parameters defining the shape of the Gaussian function were fixed at *µ* = 0 mm, axial *σ*_*a*_ = 10 mm, and radial *σ*_*r*_ = 4 mm; where axial and radial axes were parallel and perpendicular to *ω*_*v*_ respectively. For a given voxel *i* in *T*_*ab*_, the maximal value of *ϕ* across all vertices *v* was assigned:

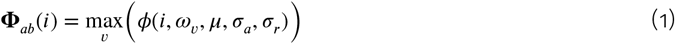

**P**_*ab*−tract_ was determined as:

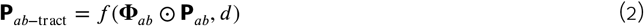

The function *f* (*g,d*) normalizes *g*, at each discrete distance *d*, to values between 0 and 1.

### Tract-specific anisotropy estimation

Having defined (or rejected) a tract *T*_*ab*_ we were next interested in extracting meaningful diffusion metrics from it, which can allow us to perform statistical inference on specific tracts. At each voxel, we thus wanted to estimate how strongly its diffusion weighed onto the orientation of the tract at that voxel. To do this, we obtained the average orientation (across all participants) of streamlines going through voxels in *T*_*ab*_ using `probtrackx` (see **Probabilistic tractography**). These average orientation images, along with the **P**_*ab*−tract_ images, were warped from standard to individual participants’ diffusion space, using the inverse transforms computed in the preprocessing step (see **Preprocessing**).

Next, for each participant, and for each voxel *j*, the fraction of the diffusion-weighted signal along the average tract orientation was estimated by fitting the following linear regression using the `statsmodels` Python library (https://www.statsmodels.org):

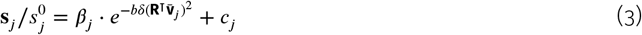

where 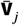 is the average streamline orientation, **R** is the *M* × 3 matrix of gradient orientation vectors (where *M* is the number of orientations, here *M* = 137), 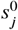 is the non-diffusion-weighted signal, **s**_*j*_ is the observed signal at each gradient orientation, *b* is the gradient strength (b-value), and *δ* is the diffusivity.

Notably, this formulation is equivalent to that presented in ***Behrens et al. (2003***, 2007), but applying only to the average orientation 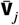. *β* _*j*_ is the regression coefficient (with *c*_*j*_ being an intercept term), and is analogous to the *f* value from the crossing fibres model; i.e., the fraction of signal contributed by 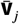. We refer to these coefficients as *tract-specific anisotropy* (TSA).

### Statistical analyses

The TSA values obtained in the preceding step were warped back into standard space and smoothed with a Gaussian kernel with a full-width at half-maximum (FWHM) of 1.5 mm. Subsequently, for each voxel in a tract, the following linear regression model was tested, for the first-order effects of *Age* and *Sex*, and their interaction:

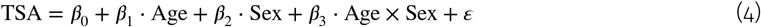

For each contrast, we next wanted to summarize the resulting t-statistics at each discrete distance along the trajectory of the tract. To do this, following an approach similar to TBSS (***Smith et al., 2006***), we computed weighted t-statistics for each voxel *j* as *t*′(*j*) = *t*(*j*) **P**_*ab*−tract_ (*j*) ^*λ*^, where *λ* determines the rate of decay (here, we set *λ* = 1.0). At each discrete distance *d* along the tract, we assigned (unweighted) *t*_*d*_ as the t-statistic corresponding to the maximal 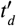.

Because anatomical properties along a tract can be assumed to form a continuous random field (i.e., neighbouring vertices have a spatial dependence), we analyzed the resulting distance-wise summary statistics as a one-dimensional random field, using the `rft1d` Python library (http://www.spm1d.org/rft1d/; ***Pataky, 2016***). Firstly, the spatial smoothness of model residuals was estimated for each tract as a FWHM value, and averaged across tracts. For each tract, this mean FWHM was used to compute *t*^*^, the critical t-value at = 0.05 for a Gaussian field, using an inverse survival function. Secondly, cluster-wise inference was performed to identify significant clusters along the tract, with a minimum cluster size of 3. Thirdly, p-values for all clusters, across all tracts, were corrected for family-wise error using a false discovery rate (FDR) threshold of 0.05. FDR was performed using the `statsmodels` Python library, with a two-stage non-negative FDR method (`fdr_tsbky`). Finally, to estimate the spatial extent of effects for a given tract, significant t-values were summed over that tract, for positive and negative t-values separately.

## Code and data sharing

All code for this procedure was written in Python 3 (Anaconda build), making system calls of native FSL packages, as indicated. The following third-party Python libraries were also used: `nibabel`, `nilearn`, and `statsmodels`. Source code for all procedures used in this study is freely available at https://github.com/neurocoglab/dwi-tracts. This includes: scripts to run FSL preprocessing and DWI analyses, using standard SGE- or SLURM-style parallel environments; a module to estimate core trajectories and TSA values; a module to fit general linear models to TSA values; and a module to plot the results of these. The output of these modules (polylines, surface meshes, graphs) can be visualised using ModelGUI software, an open source Java library available at https://github.com/neurocoglab/mgui-core. The NKI-Rockland data is freely available at www.nitrc.org/projects/fcon_1000/. Participant- and sample-wise data will be made freely available on the University of Nottingham Research Data Management Repository at https://doi.org/10.17639/nott.7102.

## Acknowledgements

We thank Saad Jbabdi for extremely helpful feedback, in particular regarding the TSA approach, as well as Marije ter Wal and Christopher Madan for their invaluable advice on various parts of the manuscript. 2D and 3D renderings for this publication were generated using ModelGUI, a free and open source software package (http://www.modelgui.org/). Parallel processing at the University of Nottingham was provided through its Augusta HPC service and the Beacon of Excellence for Precision Imaging, which provide access to high performance computing resources for the University’s neuroimaging research community.

This study was supported by the the National Institute of Mental Health (R01-MH074457), the Helmholtz Portfolio Theme “Supercomputing and Modeling for the Human Brain” and the European Union’s Horizon 2020 Research and Innovation Programme under Grant Agreement No. 4553 (HBP SGA3).

---

**Figure 5–Figure supplement 1.**
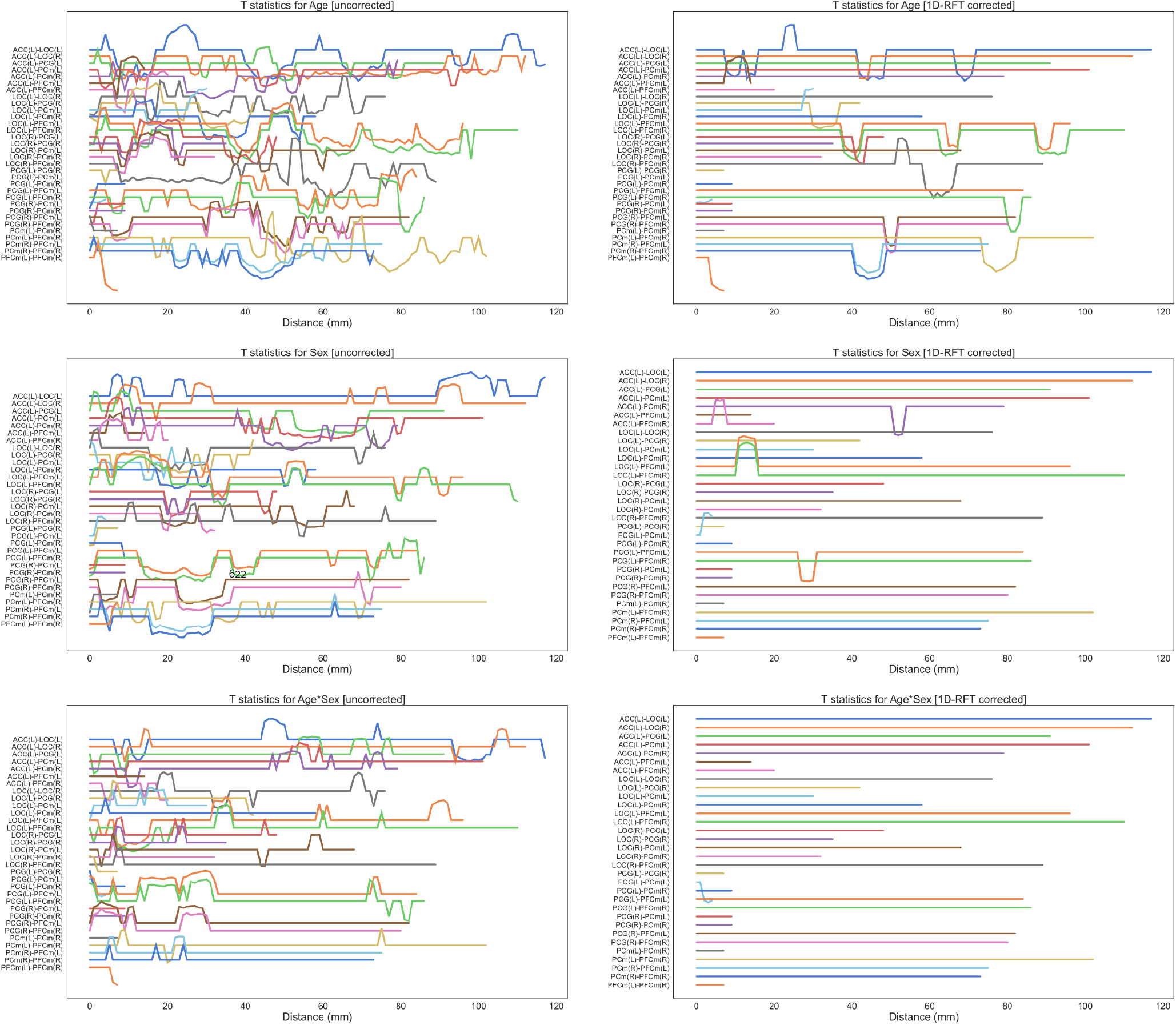
Distance traces show the t-values for *Age, Sex*, and *Age* × *Sex*, for all ROI pairs in the DMN. Plots on the left show uncorrected t-values, and plots on the right show t-values derived from one-dimensional random field theory (1D-RFT) and false-discovery rate (FDR) < 0.05. Line separation is *t* = 1.

**Figure 5–Figure supplement 2.**
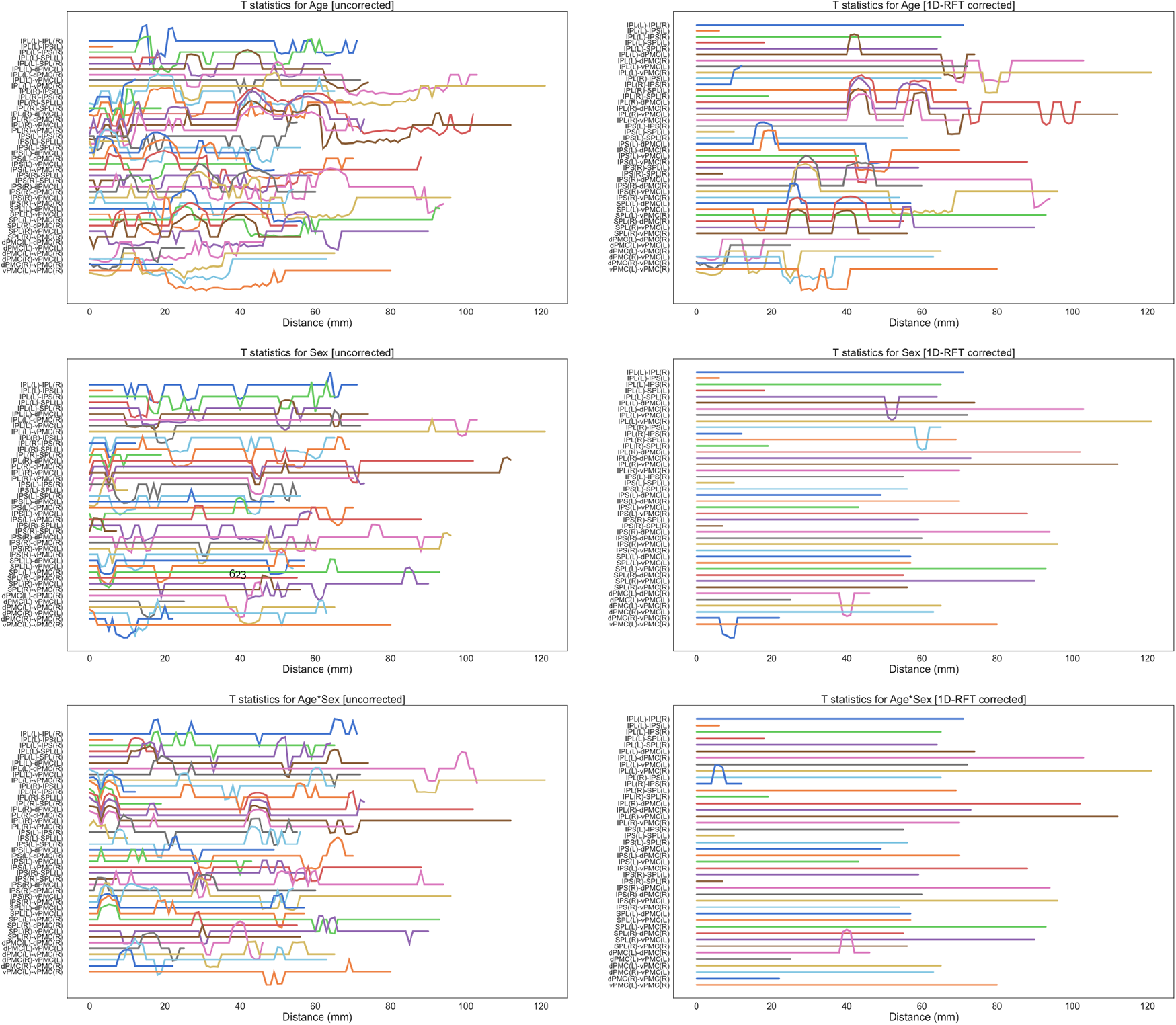
Distance traces show the t-values for *Age, Sex*, and *Age* × *Sex*, for all ROI pairs in the WWN. Plots on the left show uncorrected t-values, and plots on the right show t-values derived from one-dimensional random field theory (1D-RFT) and false-discovery rate (FDR) < 0.05. Line separation is *t* = 1.

**Figure 6–Figure supplement 1.**
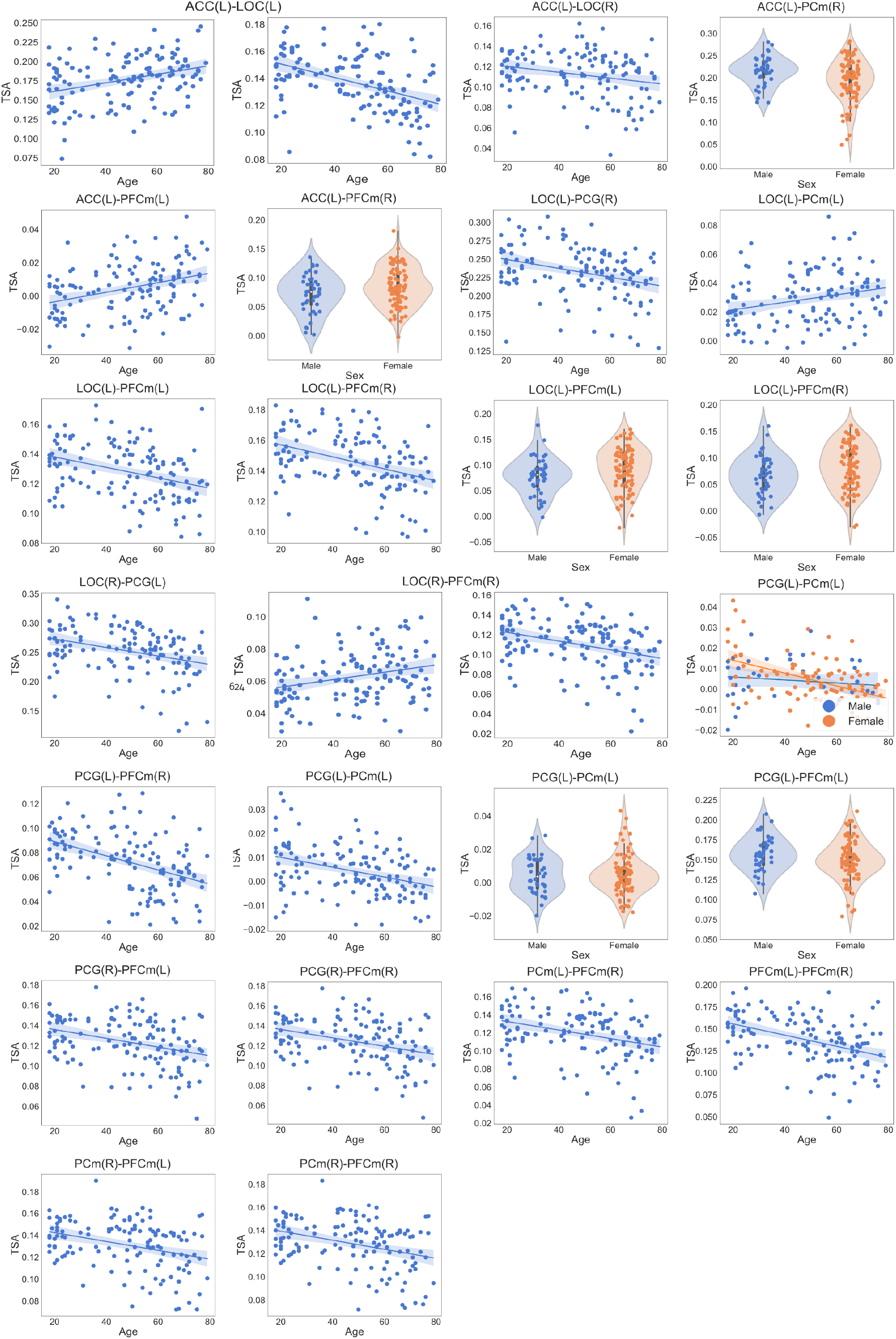
Scatter and violin plots showing significant effects for all tracts in the DMN.

**Figure 6–Figure supplement 2.**
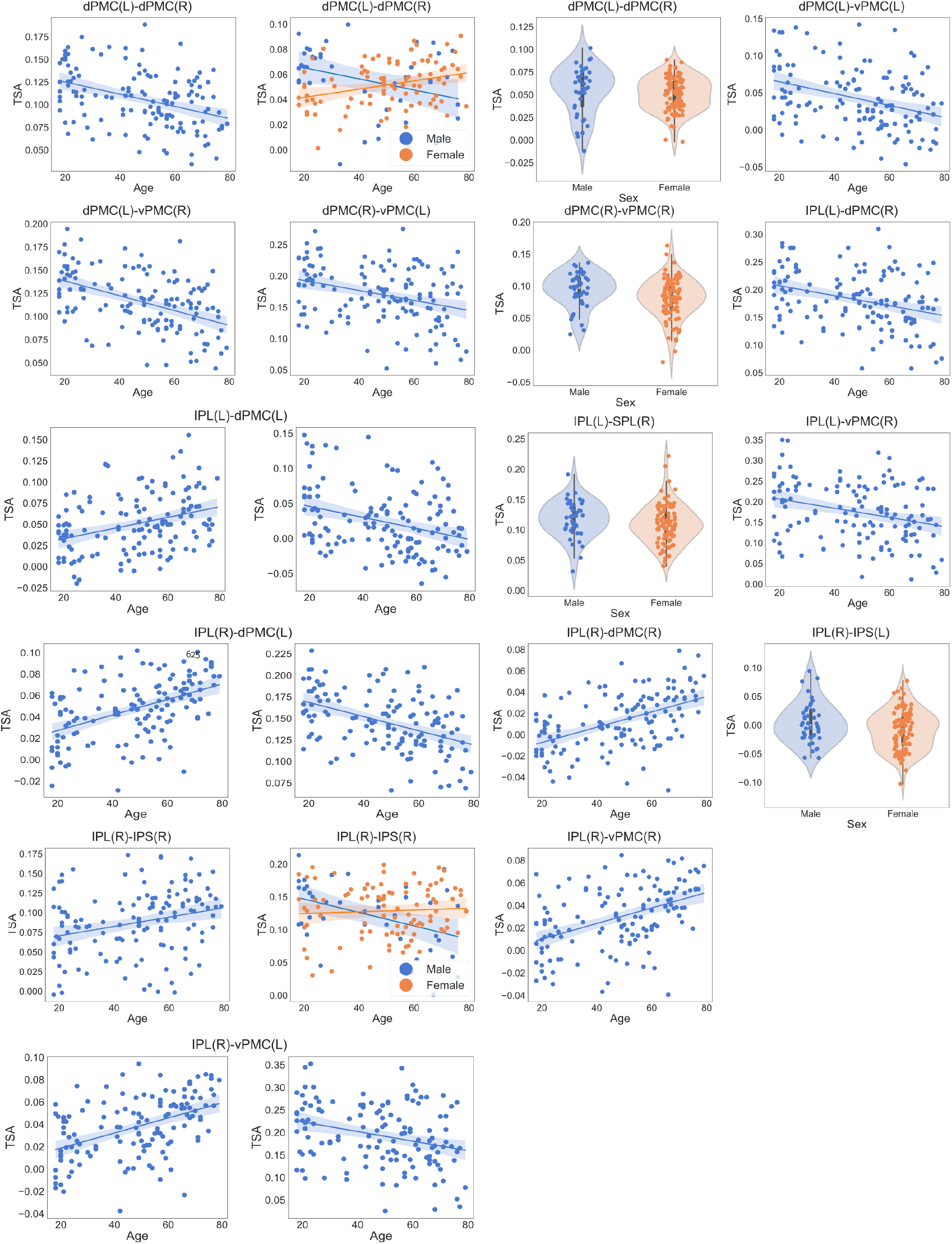
Scatter and violin plots showing significant effects for all tracts in the WWN (Part A).

**Figure 6–Figure supplement 3.**
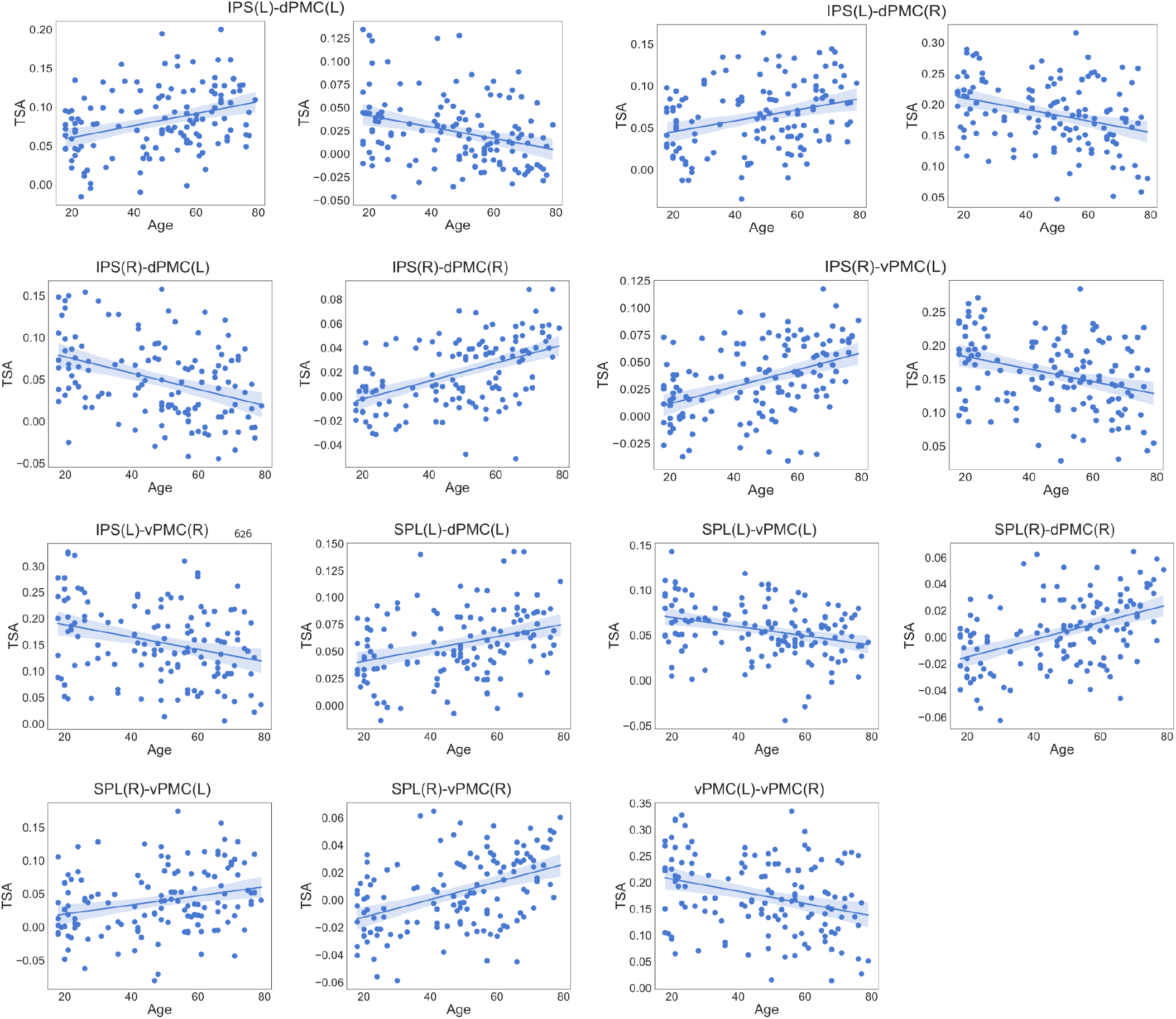
Scatter plots showing significant effects for all tracts in the WWN (Part B).

